# Perinatal Asphyxia Alters Physiological Responses and *Ex Vivo* Cardiovascular Function of Preterm Growth-Restricted Lambs

**DOI:** 10.1101/2025.05.27.656490

**Authors:** Zahrah Azman, Beth R. Piscopo, Amy E. Sutherland, Alison Thiel, Valerie A. Zahra, Yen Pham, Ilias Nitsos, Mumu Mahjabin Hossain, Atul Malhotra, Suzanne L. Miller, Kristen J. Bubb, Graeme R. Polglase, Beth J. Allison

**Author notes:** **Corresponding author:** Zahrah Azman, 27-31 Wright St, Clayton, VIC 3168, Australia. **Co-corresponding author:** Dr Beth Allison, 27-31 Wright St, Clayton, VIC 3168, Australia. These authors contributed equally (Graeme R. Polglase, Beth J. Allison).

## Abstract

**Introduction:** Fetal growth restriction (FGR) arises from chronic hypoxia and increases the risk of cardiovascular dysfunction following perinatal asphyxia, although the underlying mechanisms are unknown. We investigated whether FGR lambs have altered cardiovascular responses to perinatal asphyxia compared to control lambs, and whether impairments in α_1_ and β_1_ adrenergic receptor function underlie these responses.

**Methods:** Single or twin-bearing ewes underwent sterile fetal surgery at 89 days gestation (dGA; term=148d) to induce FGR (single umbilical artery ligation) or sham surgery (Control). At 126dGA, lambs were delivered via caesarean section, instrumented and randomised to immediate ventilation (Control_VENT_ *n*=6; FGR_VENT_ *n*=6), or asphyxia (Control_ASPHYXIA_ *n*=12; FGR_ASPHYXIA_ *n*=11) induced by umbilical cord occlusion while withholding resuscitation until diastolic blood pressure (BP) decreased to 10mmHg. Lambs were ventilated for 8 hours before baseline *ex vivo* cardiac function was assessed via Langendorff perfusion to measure left ventricular developed pressure (LVDP), heart rate (HR) and coronary perfusion pressure (CPP). *Ex vivo* α_1_ and β_1_ adrenergic responses were assessed via phenylephrine (10^−5^ to 10^−2^ mmol/L) and dobutamine (10^−7^ to 10^−4^) administration, respectively.

**Results:** FGR_ASPHYXIA_ lambs had lower BP during asphyxia (p<0.05 vs Control_ASPHYXIA_) and took longer to reach a diastolic BP of 10mmHg (14.5 ± 0.8 min vs. 19.2 ± 1.3 min; p=0.005). FGR_ASPHYXIA_ lambs had lower BP in the first 5 minutes after return of spontaneous circulation (p<0.05) due to impaired vascular contractility, with reduced Tau, dP/dt_max_ and dP/dt_min_ (p<0.03 vs Control_ASPHYXIA_). Baseline LVDP, HR and CPP were similar between groups, however FGR_ASPHYXIA_ lambs had increased LVDP responses to phenylephrine and dobutamine (p<0.05 vs Control_ASPHYXIA_), without significant changes to HR or CPP.

**Conclusion:** FGR lambs have altered physiological responses to perinatal asphyxia due to impaired vascular contractility and dysregulated cardiac α_1_ and β_1_ adrenergic receptor function, which may increase susceptibility to cardiovascular dysfunction in the neonatal period.

## Introduction

Perinatal asphyxia is the largest contributor to neonatal mortality worldwide and describes a period of severe oxygen deprivation or hypoxia in the immediate period surrounding birth^1^. Asphyxiated infants undergo a sequence of cardiovascular changes involving the redistribution of cardiac output to maintain adequate perfusion and oxygenation of critical organs such as the brain, heart and adrenal glands^2^. Increasing exposure to hypoxia leads to cardiovascular dysfunction, ultimately resulting in low cardiac output, reduced myocardial contractility secondary to poor perfusion and ischemic cardiac injury^2, 3^. Importantly, clinical and preclinical data have shown that perinatal asphyxia significantly increases the risk of long-term cardiovascular disease^4, 5^. Current resuscitation guidelines are not tailored to personalised care and therefore does not consider the influence of pre-existing perinatal complications, such as fetal growth restriction (FGR)^6^.

FGR is a pregnancy complication describing a fetus that fails to reach its biological growth potential following placental insufficiency, resulting in a state of chronic hypoxia^7–9^. FGR greatly increases the risk of cardiorespiratory complications in the neonatal period, including respiratory distress^10^, systolic and diastolic cardiac dysfunction^11–13^, and impaired autonomic control^14^. These factors likely increase the vulnerability of FGR infants experiencing perinatal asphyxia. Clinical cohort studies have demonstrated no clear consensus on whether there is a greater incidence of asphyxia in FGR infants^10, 15–17^. Regardless, FGR infants exposed to a secondary asphyxia have an increased risk of developing poorer outcomes following asphyxia, such as moderate to severe metabolic acidosis^18^. However, the physiology and mechanisms of the cardiovascular response to an asphyxia-induced acute hypoxia following chronic hypoxia resulting in FGR are poorly characterised.

In the fetus, an acute hypoxia triggers the activation of compensatory mechanisms to maintain oxygen delivery to critical organs^19^. Acute hypoxia is detected by chemoreceptors in the carotid body resulting in the redistribution of cardiac output to vital organs via the brainstem, led by an increased catecholamine production, increased sympathetic activity and peripheral vasoconstriction^19, 20^. Chronic hypoxia, as in FGR, results in a persistent redistribution of cardiac output and peripheral vasoconstriction maintained by catecholamines and adrenergic receptor signalling^19^. Several preclinical sheep studies have demonstrated that chronic hypoxia alters the cardiovascular responses to a secondary acute hypoxia, both *in utero* and in the immediate perinatal period, highlighting key deficiencies in the systems required to tolerate secondary acute hypoxia adequately^21–23^. Impairments in both α_1_ and β_1_ adrenergic signalling may therefore underlie the impeded ability of FGR neonates to defend against superimposed challenges, however these mechanisms remain unexplored in asphyxia.

The responses of preterm FGR lambs to a mild perinatal asphyxia have been previously characterised, demonstrating an apparent tolerance to asphyxia^23^. However, the mechanisms underlying this response are unknown, and it is unclear whether FGR lambs tolerate a more severe asphyxic insult. In the current study, we aimed to: (i) to investigate the ability of preterm FGR lambs to mount an adaptive cardiovascular response to severe perinatal asphyxia, and (ii) to determine whether altered cardiovascular responses are attributed to altered adrenergic receptor signalling. We hypothesised that FGR lambs will demonstrate a tolerance to severe perinatal asphyxia but will experience a compromised cardiovascular response in the recovery period due to a diminished capacity to activate myocardial α_1_ and β_1_ adrenergic receptor signalling.

## Methods

### Ethical Approval

All experimental procedures were approved by the Monash Medical Centre Animal Ethics Committee A (approval numbers MMCA 2022/08 and 2022/16) and conducted in accordance with the National Health and Medical Research Council of Australia Code of Practice for the Care and Use of Animals for Scientific Purposes. Experimental procedures were conducted in accordance with the ARRIVE (Animal Research: Reporting In Vivo Experiments) Guidelines, version 2.0.

### Animal Care and Surgical Preparation

A total of 22 singleton or twin-bearing pregnant Border-Leicester ewes underwent single umbilical artery ligation (SUAL) surgery to induce early-onset FGR, as previously described^24, 25^. Our ovine model of FGR induces early-onset placental insufficiency and asymmetrical FGR consistent with clinical definitions of FGR^24^. Ewes were monitored daily from surgery until experimentation and post-mortem.

### Asphyxia and Resuscitation

Ewes were administered intramuscular betamethasone (11.4 mg; Celestone Chronodose, Schering Plough, Australia) at 124 and 125dGA to rapidly mature critical organs such as the lung. At 126dGA (∼0.85 gestation) ewes were anaesthetised (20 mg/kg pentothal and maintained in inhaled isoflurane; 1–5% in 10/30% O_2_/N_2_O), and a caesarean section was performed to partially exteriorise the fetus from the uterus. Lambs were instrumented as previously described^23^. Briefly, the fetus was intubated with an appropriately sized cuffed endotracheal tube (3.5–4.5mm), which was clamped to prevent spontaneous breathing. The left jugular vein, brachial artery and femoral artery were catheterised. Ultrasonic flow probes (Transonic Systems) were placed around the left main pulmonary artery, left carotid artery and right femoral artery for continuous recording of blood flows. A near-infrared spectroscopy (NIRS) optode was placed on the head for measurement of regional cerebral oxygenation (crSO_2_, Foresight). A transcutaneous arterial saturation probe (Massimo) was placed around the tail to monitor peripheral oxygen saturation (SaO_2_) and heart rate. Prior to beginning experimentation, lung liquid was drained from the endotracheal tube, which was then re-clamped. Arterial catheters and flow probes were connected to a data acquisition system (Powerlab 16.35, ADInstruments) for continuous real-time physiological measurements, which were digitally recorded on LabChart Pro software (ADInstruments v1.8.3) for later offline analysis. Baseline recordings were acquired during this fetal period for two minutes before the start of the experimental protocol.

Ewes and their fetuses were randomly allocated to two groups: immediate mechanical ventilation for 8 hours (VENT) or asphyxia, resuscitation and mechanical ventilation for 8 hours (ASPHYXIA). While a larger cohort of lambs was included in the physiological analysis, only lambs that had a successful Langendorff procedure were included for all other outcomes. Therefore, the lambs included in this study were:

1. Control_VENT_ (*n*=6 for all outcomes)
2. FGR_VENT_ (*n*=6 for all outcomes)
3. Control_ASPHYXIA_ (*n*=12 for physiology; *n*=7 for other outcomes)
4. FGR_ASPHYXIA_ (*n*=11 for physiology; *n*=6 for other outcomes).

Asphyxia was induced by complete umbilical cord occlusion (UCO) whilst withholding respiratory support. The umbilical cord was cut, and the lamb was weighed and transferred to an infant radiant warmer. Asphyxia was continued until the femoral arterial end-diastolic blood pressure declined to ∼10 mmHg, which is equivalent to severe asphyxia and bradycardia (<60 bpm)^26^. Ventilation was initiated (Babylog 8000+, Dräger) in volume-guarantee mode at 7 mL/kg, maximum peak inspiratory pressure (PIP) 35 cmH_2_O, positive end-expiratory pressure (PEEP) 5 cmH_2_O, respiratory rate 60 breaths per minute, inspiratory time 0.3 s, and expiratory time 0.7 s. This ventilation strategy aligns with the normal tidal volume of spontaneously breathing lambs on continuous positive airway pressure^27^. Asynchronous chest compressions (90 compressions/min) were commenced if ventilation alone was inadequate in achieving a heart rate >100 bpm and femoral end-diastolic arterial pressure (MAP) ≥25 mmHg. Intravenous adrenaline (0.02 to 0.03 mg/kg of 1:10,000 adrenaline; AstraZeneca, Macquarie Park, NSW, Australia) was further administered if lambs failed to reach a heart rate >100 bpm and MAP >25 mmHg after 60 seconds of chest compressions and ventilation. Resuscitation was deemed successful upon return of spontaneous circulation (ROSC), which was defined as a femoral arterial end-diastolic blood pressure ≥25 mmHg and return of spontaneous pulsatile blood pressure and heart rate. Anaesthesia was maintained after ROSC through an intravenous infusion of alfaxalone (5–15 mg/kg Alfaxan in 5% glucose).

Lambs were ventilated for 8 hours following ROSC (ASPHYXIA lambs), or upon onset of ventilation (VENT lambs). Regular arterial blood samples were analysed during the ventilation period. The fraction of inspired oxygen (FiO_2_) was initially set at 1.0 and was gradually adjusted to maintain target blood gas values (PaO_2_ >40 mmHg, PaCO_2_ 35–45 mmHg, pH 7.30–7.45, SaO_2_ >85%). Ventilation parameters were adjusted during resuscitation to maintain the target blood gas values. Surfactant (100 mg/kg; Curosurf) was administered after 10 minutes of ventilation.

### Physiological Data Analysis

Physiological data were extracted and analysed in LabChart in either 10 s epochs every minute (baseline period, asphyxic period and first 15 minutes after ROSC), 30 s epochs every 5 minutes (20–60 minutes after ROSC) or 5-minute epochs every hour (1–8 hours after ROSC). As the total duration of asphyxia was not fixed, variables obtained during asphyxia were represented as quartiles relative to the duration of asphyxia for each individual animal. Femoral blood flow was corrected for body weight, while carotid and pulmonary blood flows were corrected for wet brain and lung weights, respectively. The LabChart Blood Pressure module was used to analyse vascular contractility parameters from the femoral blood pressure channel, including the exponential time constant of relaxation (Tau), the maximal rate of rise of contraction (dP/dt_max_), the maximal rate of fall of relaxation (dP/dt_min_), the slope of the isovolumetric relaxation period (IRP) and the contractility index.

### Langendorff Preparation

Immediately before postmortem, the rate of intravenous alfaxane was doubled to ensure anaesthetic overdose and heparin was administered (i.v. 5000 IU heparin sodium) to prevent coagulation of the coronary arteries. A cervical dislocation was conducted after confirming the absence of pain reflexes, and the chest cavity was opened to access the heart. The aorta was clamped, and the heart was swiftly excised and placed in ice-cold, oxygenated Krebs solution. Briefly, Krebs solution contained (in mM) 120 NaCl, 4.7 KCl, 1.2 MgSO_4_·7H_2_O, 1.2 KH_2_PO_4_, 25 NaHCO_3_, 10 glucose, and 1.3 CaCl_2_·H_2_O^28^. The heart was mounted on a Langendorff apparatus via the aorta and constantly perfused with Krebs solution at a constant flow rate of 40 mL/min. A saline-filled latex balloon connected to a catheter was inserted into the left ventricle and set to a diastolic pressure of 5 mmHg to measure left ventricular developed pressure (LVDP). Isolated hearts were equilibrated for 10 minutes before baseline functional recordings were acquired. Measurements of baseline function included coronary perfusion pressure (CPP; measured via pressure ejected from the aorta), heart rate, maximal, minimal, and mean LVDP, maximal rate of rise of left ventricular contraction (dP/dt_max_), and maximal rate of fall of left ventricular relaxation (dP/dt_min_). Cardiac responses to 1) β_1_-adrenergic receptor stimulation via administration of dobutamine (10^−7^ to 10^−4^ mmol/L); 2) nitric oxide donor via administration of glyceryl trinitrate (GTN; 10^−2^ mmol/L); 3) α_1_-adrenergic receptor stimulation via administration of phenylephrine (10^−5^ to 10^−2^ mmol/L) were determined. The heart was allowed to return to baseline function following the completion of each drug curve. At the end of the Langendorff protocol, hearts were dismounted and transversely sliced into 1-cm-thick sections, with the most superior section of the ventricles incubated in 1% 2,3,5-triphenyltetrahydrozolium chloride (10 mg/mL dissolved in PBS) for 15 min at 37°C to determine infarct size. Segments of left ventricle were either snap-frozen for RT-qPCR or fixed with 10% neutral-buffered formalin for histological analysis.

### RNA extraction and qPCR

Total RNA was extracted from 20–30 mg of frozen left ventricular myocardium (RNeasy Mini Kit, Qiagen) before 50 ng/µL of RNA was reverse-transcribed into cDNA (SuperScript III Reverse Transcriptase, Thermo Fisher). Relative mRNA levels of ɑ_1_ adrenergic receptor (*ɑ_1_-AR*), β_1_ adrenergic receptor (*β_1_-AR*), atrial natriuretic peptide (*NPPA*), B-type natriuretic peptide (*NPPB*) and sarcoplasmic/endoplasmic reticulum calcium ATPase 2 (*SERCA/ATP2A2*) (Supplementary Table 1) were measured by RT-qPCR (QuantStudio 6 Real-Time PCR system, ThermoFisher, USA). Genes were quantified and normalised to the housekeeping gene 18S using the cycle threshold (ΔC_T_) method. The expression of all genes was expressed relative to the mean of the Control_VENT_ group.

### Histological and immunohistochemical analyses

Left ventricle sections (5 µm thick) were stained with Masson’s trichrome^29^ to evaluate perivascular and interstitial fibrosis. Ki67 and α_1_-adrenergic receptor immunohistochemistry were performed to identify proliferative cells and α_1_-adrenergic receptor populations (Supplementary Table 2). Negative control slides confirmed the specificity of Ki67 or α_1_-adrenergic receptor staining. Qualitative assessment of the number of Ki67-positive cells, the area of vascular α_1_-adrenergic receptor staining, and interstitial and perivascular fibrosis was completed by an assessor (ZA) blinded to the experimental groups on coded slides.

### Statistical analysis

Data are expressed as the mean ± standard error of the mean (SEM) and analysed via GraphPad Prism software (GraphPad Prism 10). Data were assessed for normality using a Shapiro-Wilk test prior to further statistical analysis. Postmortem characteristics, baseline heart function, histology and molecular data were analysed via a two-way analysis of variance (ANOVA). Lamb physiology and blood gas measurements were analysed via a three-way repeated measures ANOVA with time, fetal growth (FGR or control) and exposure to asphyxia as fixed variables. *Ex vivo* cardiac responses to drug administration were analysed via a three-way ANOVA with fetal growth (FGR or control), exposure to asphyxia, and drug doses as fixed variables. A multiple comparisons uncorrected Fisher’s least significant difference test was conducted to isolate differences in significant interactions between the main factors. Comparisons were deemed statistically significant at a p-value <0.05.

## Results

### Lamb characteristics, asphyxia characteristics and blood gases

Postmortem lamb characteristics, asphyxia characteristics, baseline blood gases and end-UCO blood gases are described in Table 1. Body weight was significantly lower in FGR lambs compared to control lambs (p=0.001). Asymmetric growth restriction was evident with a significantly increased brain-to-body weight ratio (p=0.004) and similar heart-to-body weight ratio. Sex and birth order were not different between groups.

**Table 1.**
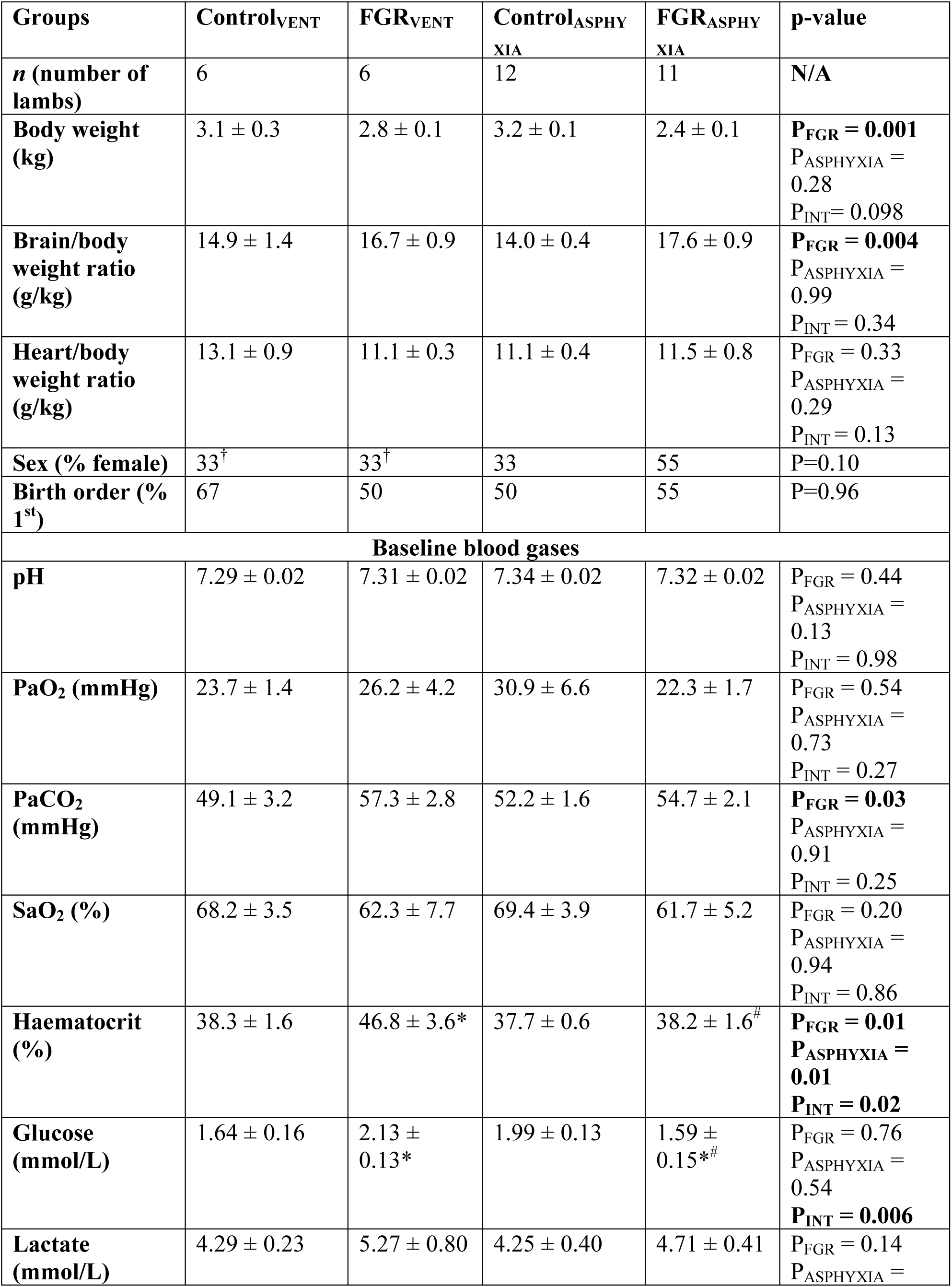
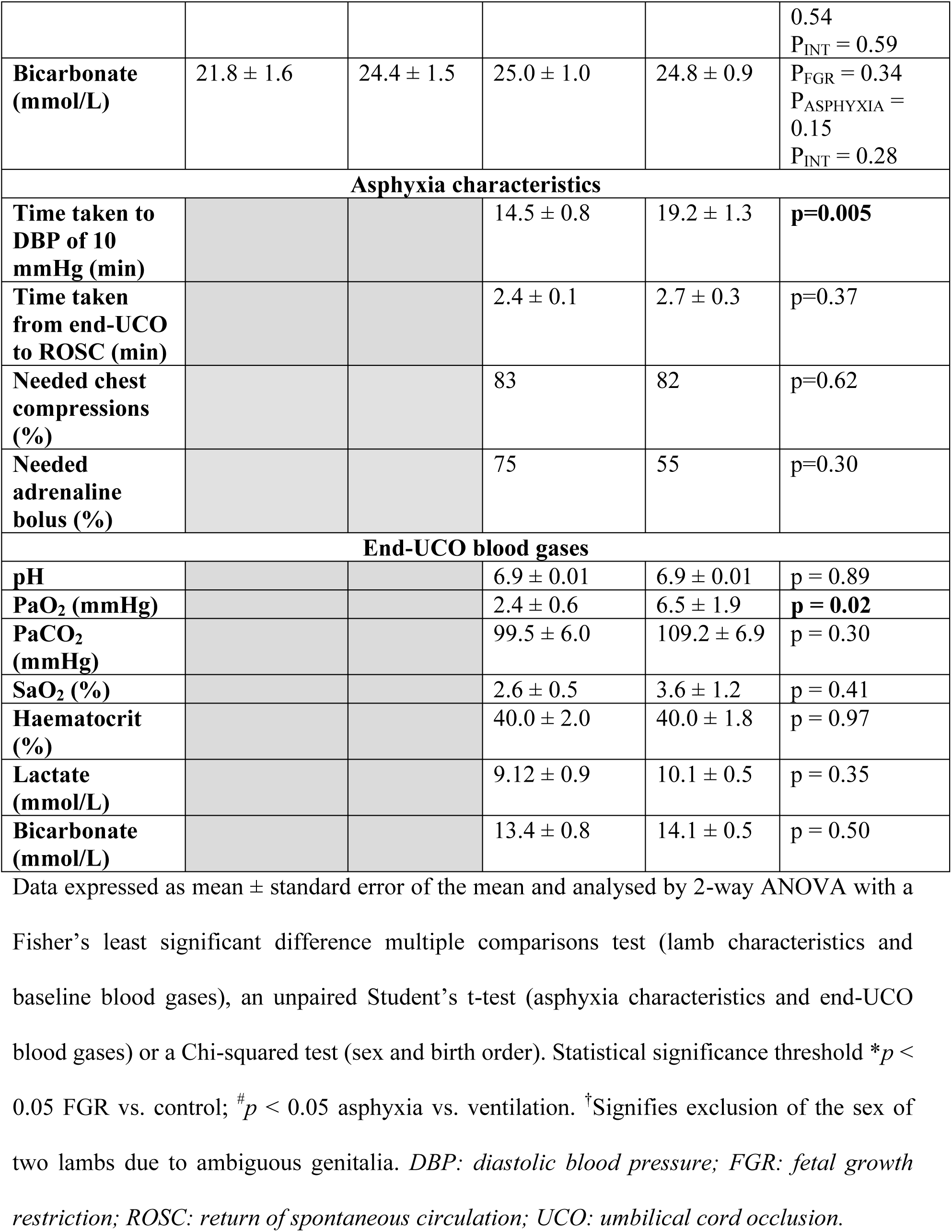
Lamb characteristics, asphyxia characteristics and blood gases.

FGR groups were hypercapnic at baseline compared to control groups (p=0.03). FGR_VENT_ lambs had significantly higher haematocrit levels than Control_VENT_ lambs (p=0.02), while FGR_ASPHYXIA_ lambs had significantly lower haematocrit levels than FGR_VENT_ lambs (p=0.009). Baseline glucose levels of FGR_VENT_ lambs were significantly higher than Control_VENT_ lambs (p=0.047), while FGR_ASPHYXIA_ lambs had significantly lower glucose levels compared to FGR_VENT_ (p=0.02) and Control_ASPHYXIA_ (p=0.04) lambs. No other differences in baseline arterial blood gas parameters were evident.

FGR_ASPHYXIA_ lambs took significantly longer to reach an end-diastolic blood pressure of 10 mmHg compared to Control_ASPHYXIA_ lambs (14.5 ± 0.8 min vs. 19.2 ± 1.3 min; p=0.005). However, the time taken to achieve ROSC did not differ between groups. The requirement for resuscitation interventions (chest compressions and adrenaline boluses) were not different between groups.

At the end of the asphyxia period, arterial pH, PaCO_2_, SaO_2_, haematocrit, lactate, and bicarbonate were not different between Control_ASPHYXIA_ and FGR_ASPHYXIA_ lambs. However, Control_ASPHYXIA_ lambs had significantly lower PaO_2_ at the end of the asphyxia period compared to FGR_ASPHYXIA_ lambs (p=0.02).

### Physiology during asphyxia

Mean arterial blood pressure was significantly lower in FGR_ASPHYXIA_ lambs at 50% asphyxia compared to Control_ASPHYXIA_ lambs (p=0.02; Fig. 1A). Peak systolic and end diastolic blood pressures were significantly lower in FGR_ASPHYXIA_ lambs compared to Control_ASPHYXIA_ lambs between 50%–75% of the asphyxic period (p<0.05; Fig. 1B&C). FGR_ASPHYXIA_ lambs had significantly lower pulse pressures compared to Control_ASPHYXIA_ lambs during asphyxia (p=0.047; Fig. 1D). Mean heart rate was not different between groups during asphyxia (data not shown).

**Figure 1.**
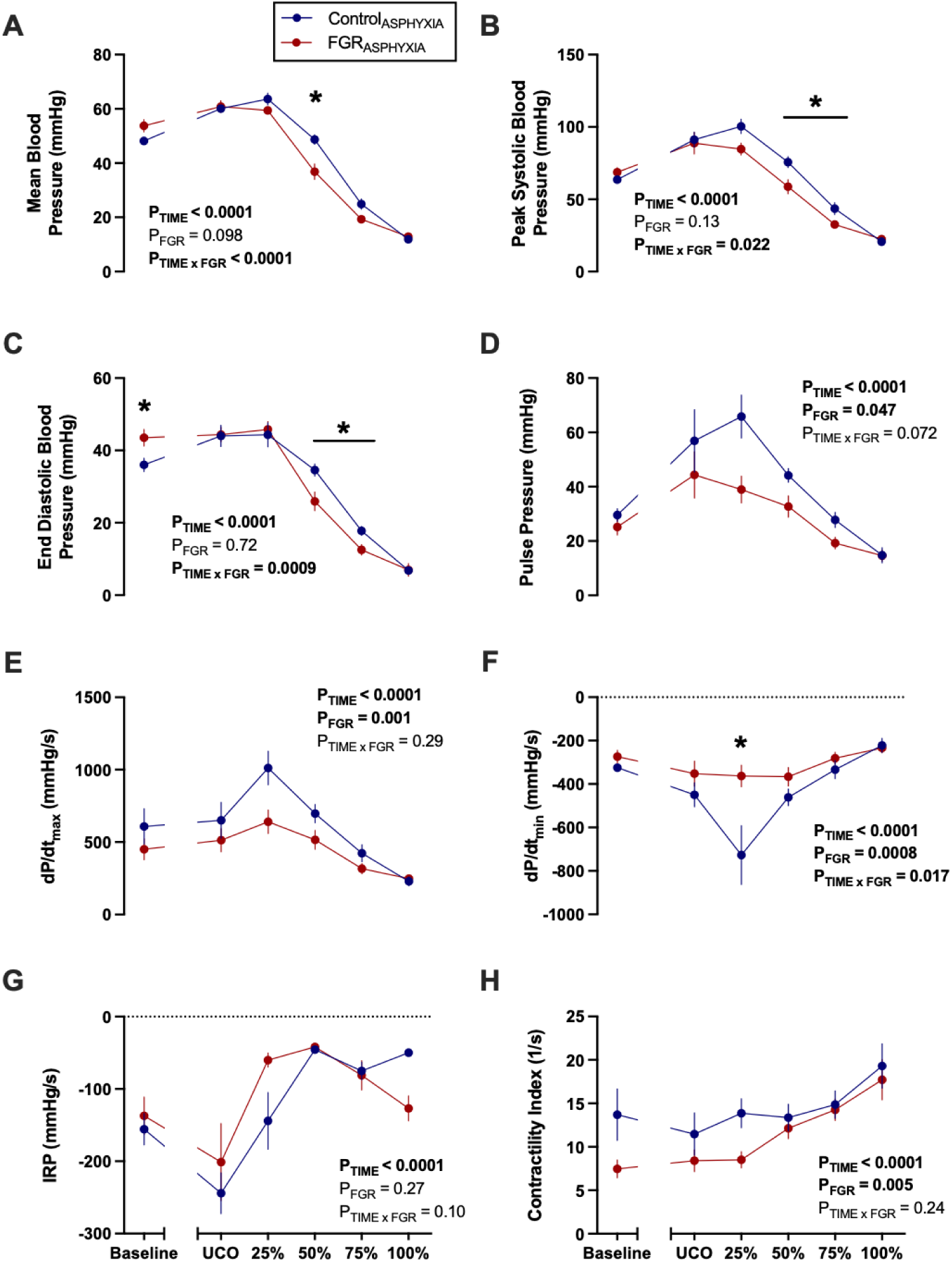
Blood pressure and vascular contractility during asphyxia. Data presented as mean ± standard error of the mean of (**A**) mean arterial blood pressure, (**B**) peak systolic blood pressure, (**C**) end diastolic blood pressure, (**D**) pulse pressure, (**E**) dP/dt_max_, (**F**) dP/dt_min_, (**G**) isovolumetric relaxation period (IRP) and (**H**) contractility index. Data are presented at baseline, umbilical cord occlusion (UCO), and at quartiles relative to the total duration of asphyxia (25, 50, 75 and 100%). Groups are asphyxiated control (Control_ASPHYXIA_, n=12), and asphyxiated FGR (FGR_ASPHYXIA_, n=11) lambs. Data was analysed via a repeated measure mixed effects analysis with Fisher’s least significant difference multiple comparisons test. \**p*<0.05.

Vascular contractility parameters were also derived from the arterial pressure signal (Fig. 1E-H). FGR_ASPHYXIA_ lambs had a significantly lower dP/dt_max_, dP/dt_min_, and contractility index compared to Control_ASPHYXIA_ lambs throughout the asphyxic period (p<0.05; Fig 1E,F,H). Control_ASPHYXIA_ lambs had a steeper decline in dP/dt_min_ (p<0.0001; Fig. 1F) compared to FGR_ASPHYXIA_ lambs at 25% asphyxia.

Carotid and pulmonary blood flows were not different between Control_ASPHYXIA_ and FGR_ASPHYXIA_ lambs during asphyxia (Fig. 2A&B). However, femoral blood flow was significantly lower in FGR_ASPHYXIA_ than Control_ASPHYXIA_ at baseline and immediately after UCO (p<0.05; Fig. 2C). FGR_ASPHYXIA_ lambs had a higher carotid to femoral blood flow ratio during asphyxia (p=0.03; Fig. 2D). FGR_ASPHYXIA_ lambs also had a significantly lower regional cerebral oxygenation during asphyxia compared to Control_ASPHYXIA_ lambs (p=0.02; Fig. 2E).

**Figure 2.**
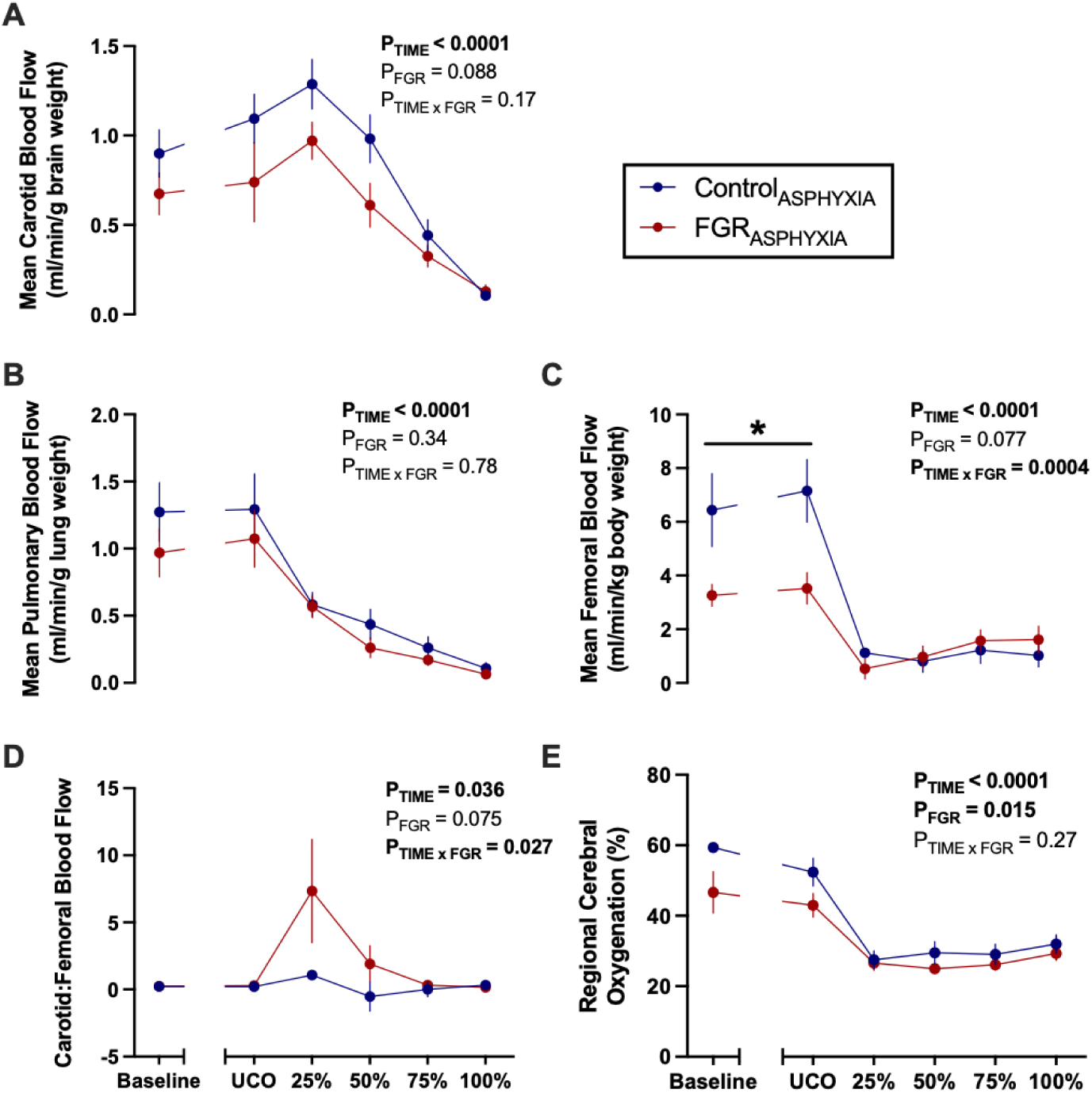
Blood flows and regional cerebral oxygenation during asphyxia. Data presented as mean ± standard error of the mean of (**A**) mean carotid blood flow (corrected for brain weight), (**B**) mean pulmonary blood flow (corrected for lung weight), (**C**) mean femoral blood flow (corrected for body weight), (**D**) ratio of carotid to femoral blood flow and (**E**) mean regional cerebral oxygenation. Data are presented at baseline, umbilical cord occlusion (UCO), and at quartiles relative to the total duration of asphyxia (25, 50, 75 and 100%). Groups are asphyxiated control (Control_ASPHYXIA_, n=12) and asphyxiated FGR (FGR_ASPHYXIA_, n=11) lambs. Data analysed via repeated measure mixed effects analysis with Fisher’s least significant difference multiple comparisons test. \**p*<0.05.

### Blood gases during ventilation

Arterial pH, PaCO_2_, SaO_2_ and glucose levels were similar between groups during ventilation (Supplementary Fig. 1A&B, D&E). Control_ASPHYXIA_ lambs had significantly higher PaO_2_ compared to Control_VENT_ animals at 10 and 15 minutes after ROSC (p<0.05; Supplementary Fig. 1C). FGR_ASPHYXIA_ lambs had a significantly lower PaO_2_ than FGR_VENT_ lambs at 10 minutes, but this rebounded resulting in a significantly higher PaO_2_ than FGR_VENT_ lambs at 15 minutes (p<0.05; Supplementary Fig. 1C). Lactate concentrations were significantly higher in Control_ASPHYXIA_ lambs compared to Control_VENT_ lambs from 5 minutes to 2 hours after ROSC (p<0.05; Supplementary Fig. 1F). Lactate concentrations were also significantly higher in FGR_ASPHYXIA_ lambs compared to FGR_VENT_ animals between 5 minutes–2 hours and at 4 hours after ROSC (p<0.05; Supplementary Fig. 1F). FGR_ASPHYXIA_ lambs also had significantly higher lactate levels at 8 hours after ROSC compared to Control_ASPHYXIA_ lambs (p=0.03; Supplementary Fig. 1F).

### Physiology during ventilation

#### Blood pressure and vascular contractility

Blood pressure, heart rate and pulse pressure after ROSC and during mechanical ventilation are shown in Figure 3. At ROSC, FGR_ASPHYXIA_ lambs had a significantly lower mean arterial blood pressure than Control_ASPHYXIA_ (p=0.048) and FGR_VENT_ lambs (p=0.02; Fig. 3A). Control_ASPHYXIA_ lambs had a significantly higher mean arterial blood pressure compared to Control_VENT_ lambs for the first 20 minutes after ROSC (p<0.05; Fig. 3A) and compared to FGR_ASPHYXIA_ lambs for the first 5 minutes after ROSC (p<0.05; Fig. 3A). However, the mean arterial blood pressure of FGR_ASPHYXIA_ lambs began to rise after 5 minutes and was significantly higher than FGR_VENT_ lambs between 9 to 15 minutes after ROSC (p<0.05; Fig. 3A). This rebound relative hypertension was transient in FGR_ASPHYXIA_ lambs and returned to a significantly lower mean arterial blood pressure compared to Control_ASPHYXIA_ lambs by 20 minutes (Fig. 3A; p=0.049).

**Figure 3.**
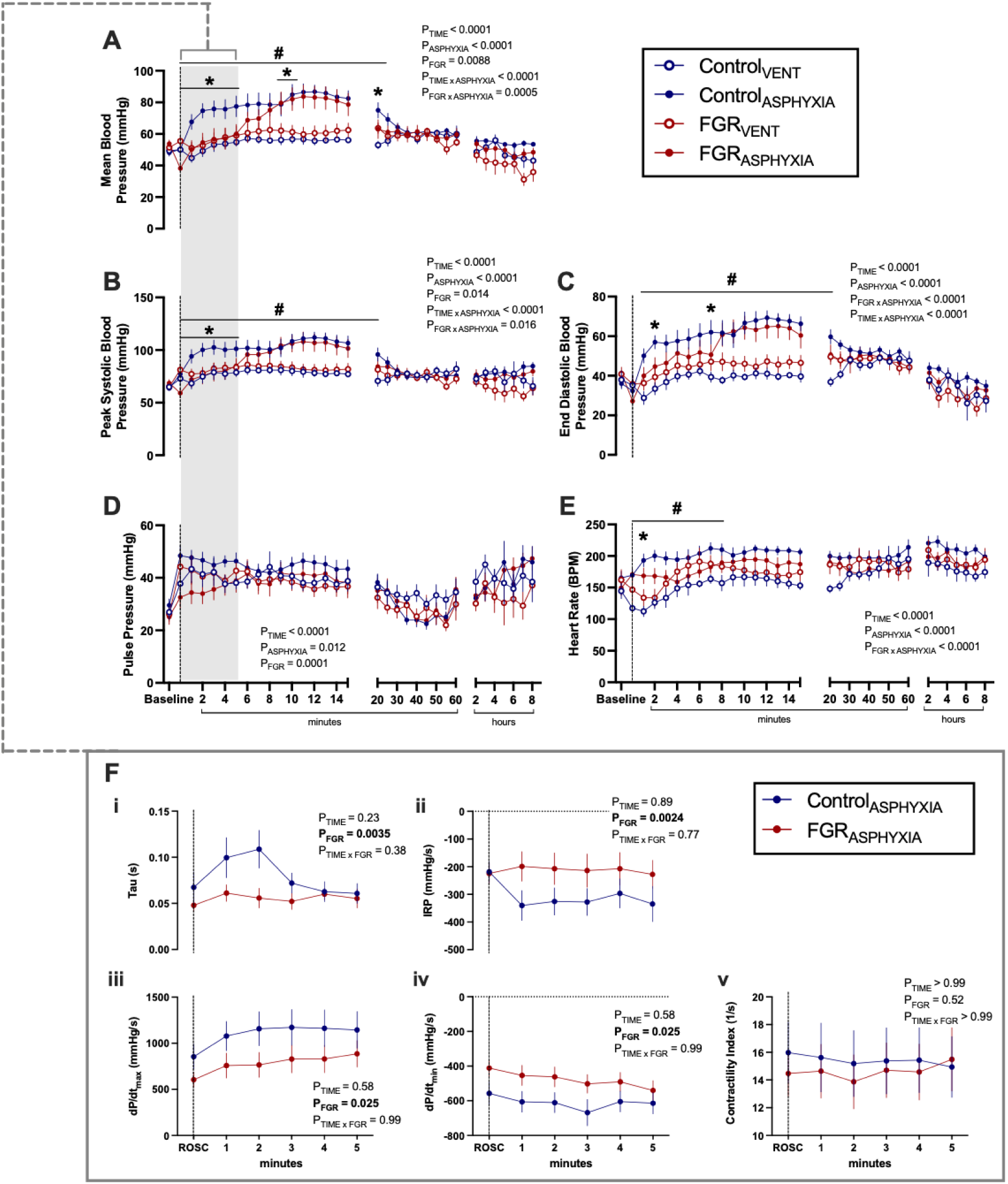
Blood pressure and vascular contractility during ventilation. Data presented as mean ± standard error of the mean of (**A**) mean blood pressure, (**B**) peak systolic blood pressure, (**C**) end diastolic blood pressure, (**D**) pulse pressure, (**E**) mean heart rate, and (**F**) Blood pressure-derived vascular contractility in the first 5 minutes after ROSC, including (**i**) Tau, (**ii**) isovolumetric relaxation period (IRP), (**iii**) dP/dt_max_, (**iv**) dP/dt_min_, and (**v**) contractility index. Data are presented at baseline, at ROSC (┄ dotted line), and during 8 hours of post-recovery ventilation. Groups are ventilated control (Control_VENT_, n=6), ventilated FGR (FGR_VENT_, n=6), asphyxiated control (Control_ASPHYXIA_, n=12), and asphyxiated FGR (FGR_ASPHYXIA_, n=11) lambs. Data analysed by a repeated measure mixed effects analysis with a Fisher’s least significant difference multiple comparisons test. \**p* < 0.05 FGR vs. control; ^#^*p* < 0.05 asphyxia vs. ventilation.

At ROSC, the systolic blood pressure of FGR_ASPHYXIA_ lambs was significantly lower than FGR_VENT_ (p=0.01) and Control_ASPHYXIA_ lambs (p=0.02; Fig. 3B). Systolic blood pressure remained significantly higher in Control_ASPHYXIA_ lambs compared to FGR_ASPHYXIA_ lambs for the first 10 minutes after ROSC (p<0.05; Fig. 3B) and compared to Control_VENT_ lambs for the first 20 minutes after ROSC (p<0.05; Fig. 3B). The increase in systolic blood pressure was delayed in FGR_ASPHYXIA_ lambs and was only demonstrated after 10 minutes, with significantly higher pressures compared to FGR_VENT_ lambs between 10 to 15 minutes after ROSC (p<0.05; Fig. 3B).

End diastolic blood pressures were similar between groups at ROSC. Between 1 to 25 minutes, Control_ASPHYXIA_ lambs had significantly higher end diastolic blood pressures compared to Control_VENT_ lambs (p<0.05; Fig. 3C) and at 2 minutes and 7 minutes after ROSC compared to FGR_ASPHYXIA_ lambs (p<0.05; Fig. 3C). FGR_ASPHYXIA_ lambs also had significantly lower end diastolic blood pressure compared to FGR_VENT_ lambs between 8 to 15 minutes (p<0.05; Fig. 3C).

Mean, systolic and diastolic blood pressure levels remained similar between groups from 30 minutes until the end of the ventilation period.

The heart rate of Control_ASPHYXIA_ lambs was significantly higher than Control_VENT_ lambs at ROSC and for the first 8 minutes after ROSC (p<0.05; Fig. 3D) and compared to FGR_ASPHYXIA_ lambs between 2 and 7 minutes after ROSC (p<0.05; Fig. 3D). FGR_ASPHYXIA_ lambs had a significantly higher heart rate compared to FGR_VENT_ lambs 1 minute after ROSC (p=0.046; Fig. 3D). Pulse pressures were not different between groups at ROSC or during the ventilation period (Fig. 3E).

The vascular contractility of Control_ASPHYXIA_ and FGR_ASPHYXIA_ lambs in the first 5 minutes after ROSC was further investigated via the arterial blood pressure waveform (Fig. 3F). In the first 5 minutes after ROSC, Tau, IRP, dP/dt_max_ and dP/dt_min_ were significantly lower in FGR_ASPHYXIA_ lambs compared to Control_ASPHYXIA_ lambs (p<0.05; Fig. 3F[i–iv]). However, the contractility index was not different between groups during this period (Fig. 3F[v]).

#### Blood flows and regional cerebral oxygenation

Mean carotid blood flow was higher in FGR_ASPHYXIA_ compared to FGR_VENT_ lambs between 11–12 minutes after ROSC (p<0.05; Fig. 4A). Mean carotid blood flow was also higher in Control_ASPHYXIA_ compared to Control_VENT_ lambs at 20 and 25 minutes (p<0.05; Fig. 4A) and compared to FGR_ASPHYXIA_ lambs at 25 minutes (p=0.008; Fig. 4A).

**Figure 4.**
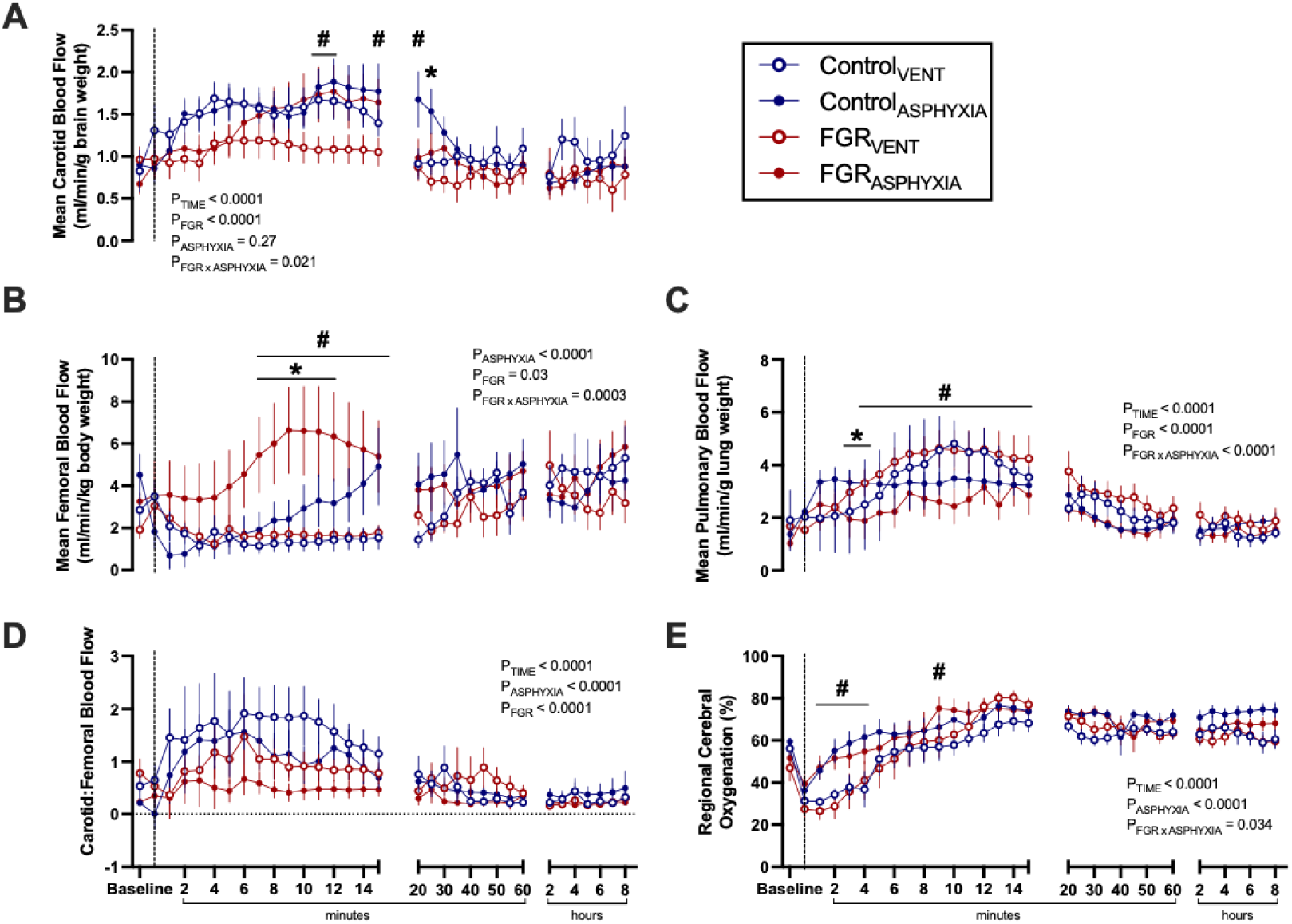
Blood flows and regional cerebral oxygenation during ventilation. Data presented as mean ± standard error of the mean of (**A**) mean carotid blood flow (corrected for brain weight), (**B**) mean femoral blood flow (corrected for body weight), (**C**) mean pulmonary blood flow (corrected for lung weight), (**D**) ratio of carotid to femoral blood flow and (**E**) mean regional cerebral oxygenation. Data are presented at baseline, at return of spontaneous circulation ( ┄ dotted line), and during 8 hours of post-recovery ventilation. Groups are ventilated control (Control_VENT_, n=6), ventilated FGR (FGR_VENT_, n=6), asphyxiated control (Control_ASPHYXIA_, n=12), and asphyxiated FGR (FGR_ASPHYXIA_, n=11) lambs. Data analysed via a repeated measure mixed effects analysis with a Fisher’s least significant difference multiple comparisons test. \**p* < 0.05 FGR vs. control; ^#^*p* < 0.05 asphyxia vs. ventilation.

Mean femoral blood flow was higher in FGR_ASPHYXIA_ compared to Control_ASPHYXIA_ lambs between 7–12 minutes after ROSC (p<0.05; Fig. 4B) and compared to FGR_VENT_ lambs between 7–15 minutes after ROSC (p<0.05; Fig. 4B). The ratio of carotid to femoral blood flow was not different between groups after ROSC (Fig. 4D).

Mean pulmonary blood flow was lower in FGR_ASPHYXIA_ lambs compared to Control_ASPHYXIA_ lambs between 3–4 minutes after ROSC (p<0.05; Fig. 4C) and compared to FGR_VENT_ lambs between 4–15 minutes after ROSC (p<0.05; Fig. 4C).

Regional cerebral oxygenation was higher in FGR_ASPHYXIA_ lambs compared to FGR_VENT_ lambs from 1–3 minutes and at 9 minutes after ROSC (p<0.05; Fig. 4E). Regional cerebral oxygenation was also higher in Control_ASPHYXIA_ lambs compared to Control_VENT_ lambs between 2–4 minutes after ROSC (p<0.05; Fig. 4E).

### Langendorff perfusion

Isolated lamb hearts were assessed via a Langendorff perfusion to understand the isolated role of cardiac function in response to a secondary asphyxia. Basal isolated cardiac function, including coronary perfusion pressure, heart rate, mean and maximal LVDP amplitude, and the maximal rate of rise of contraction (dP/dt_max_) and relaxation (dP/dt_min_) were similar between groups (Supplementary Fig. 2).

Hearts from ventilated lambs experienced a greater increase in coronary perfusion pressure in response to dobutamine compared to hearts from asphyxiated lambs (p=0.006; Fig. 5A). FGR_ASPHYXIA_ hearts had a greater increase in mean, amplitude and maximal LVDP following a 10^−7^ dose of dobutamine compared to Control_ASPHYXIA_ and FGR_VENT_ hearts (p<0.05; Fig. 5B–E). FGR_ASPHYXIA_ hearts also had a greater increase in maximal LVDP compared to FGR_VENT_ hearts following a 10^−4^ dose of dobutamine (p=0.049; Fig. 5E). Asphyxiated hearts also demonstrated a larger increase in dP/dt_max_ and dP/dt_min_ following dobutamine (p<0.05; Fig. 5F&G). Heart rate responses following dobutamine were not different between groups (Fig. 5B).

**Figure 5.**
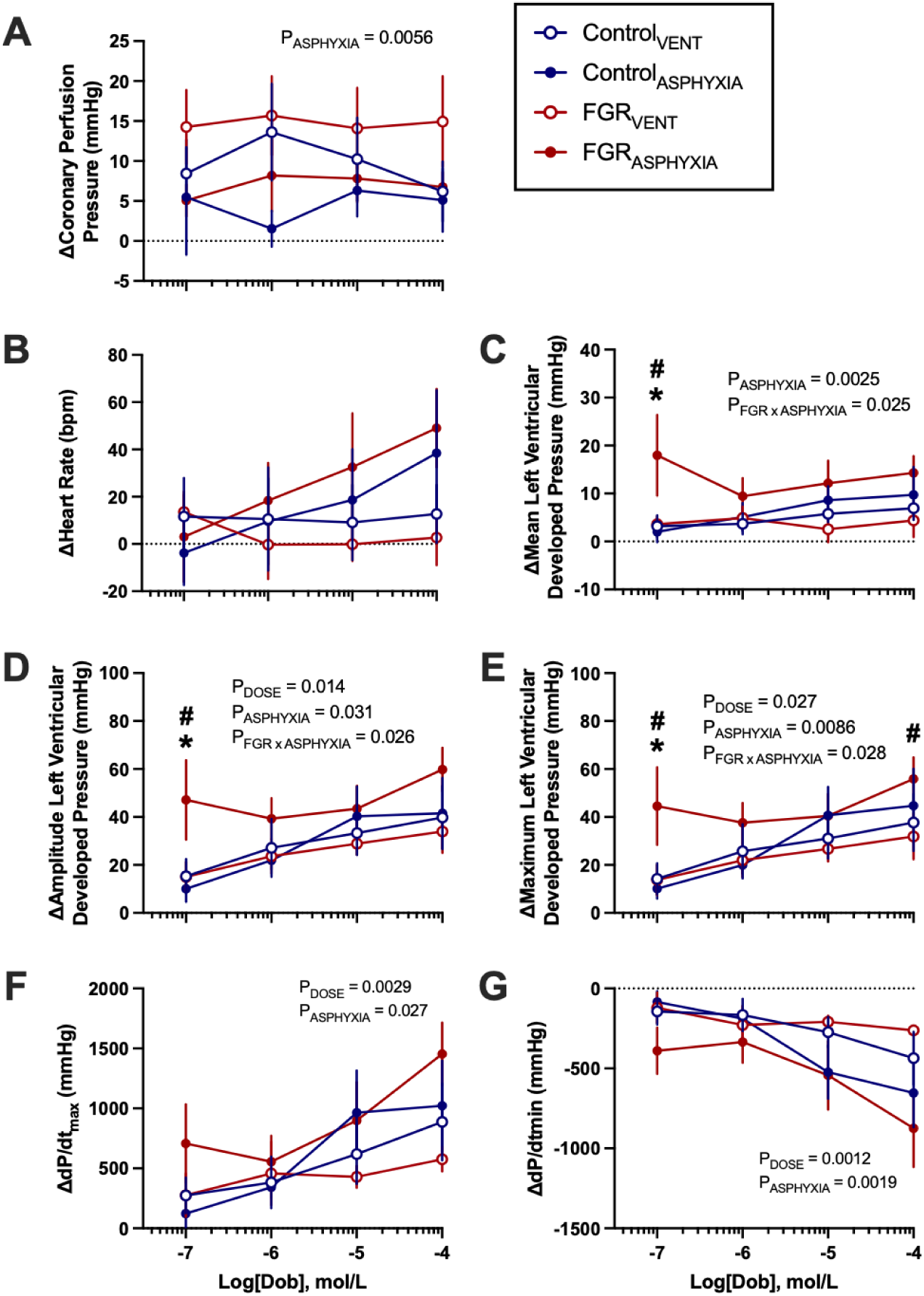
Ex vivo cardiac responses to dobutamine. Data presented as mean ± standard error of the mean change in (**A**) coronary perfusion pressure, (**B**) heart rate, (**C**) mean left ventricular developed pressure, (**D**) amplitude of left ventricular developed pressure, (**E**) maximum left ventricular developed pressure, (**F**) maximal left ventricular contractility (dP/dt_max_) and (**G**) minimal left ventricular contractility (dP/dt_min_) in response to increasing concentrations of dobutamine. Groups are ventilated control (Control_VENT_, n=6), ventilated FGR (FGR_VENT_, n=6), asphyxiated control (Control_ASPHYXIA_, n=7), and asphyxiated FGR (FGR_ASPHYXIA_, n=6) lambs. Data analysed via mixed effects analysis with uncorrected Fisher’s least significance multiple comparisons test; statistical significance threshold *p*<0.05. \**p* < 0.05 FGR vs. control; ^#^*p* < 0.05 asphyxia vs. ventilation.

Hearts from FGR_ASPHYXIA_ animals showed greater increases in mean LVDP, dP/dt_max_ and dP/dt_min_ compared to FGR_VENT_ animals in response to 10^−4^ to 10^−2^ doses of phenylephrine (p<0.05; Fig. 6C, F&G). FGR_ASPHYXIA_ animals also had greater increases in mean LVDP compared to Control_ASPHYXIA_ animals in response to 10^−3^ and 10^−2^ doses of phenylephrine (p<0.05; Fig. 6C). FGR_ASPHYXIA_ hearts also demonstrated greater increases in LVDP amplitude (10^−4^ to 10^−2^ doses) and maximal LVDP (10^−4^ to 10^−3^ doses) compared to Control_ASPHYXIA_ hearts in response to phenylephrine administration (p<0.05; Fig. 6D&E). FGR_ASPHYXIA_ hearts also had a greater amplitude and maximal LVDP response compared to FGR_VENT_ hearts following administration of the 10^−4^ dose of phenylephrine (p<0.05; Fig. 6D&E). FGR_ASPHYXIA_ hearts experienced greater increases in dP/dt_max_ (10^−4^ to 10^−2^ dose) and dP/dt_min_ (10^−3^ dose) in response to phenylephrine compared to Control_ASPHYXIA_ animals (p<0.05; Fig 6F&G). Heart rate responses to phenylephrine were not different between groups (Fig. 6B). Isolated cardiac responses to glyceryl trinitrate, a nitric oxide donor, were not different between groups (Supplementary Fig. 3).

**Figure 6.**
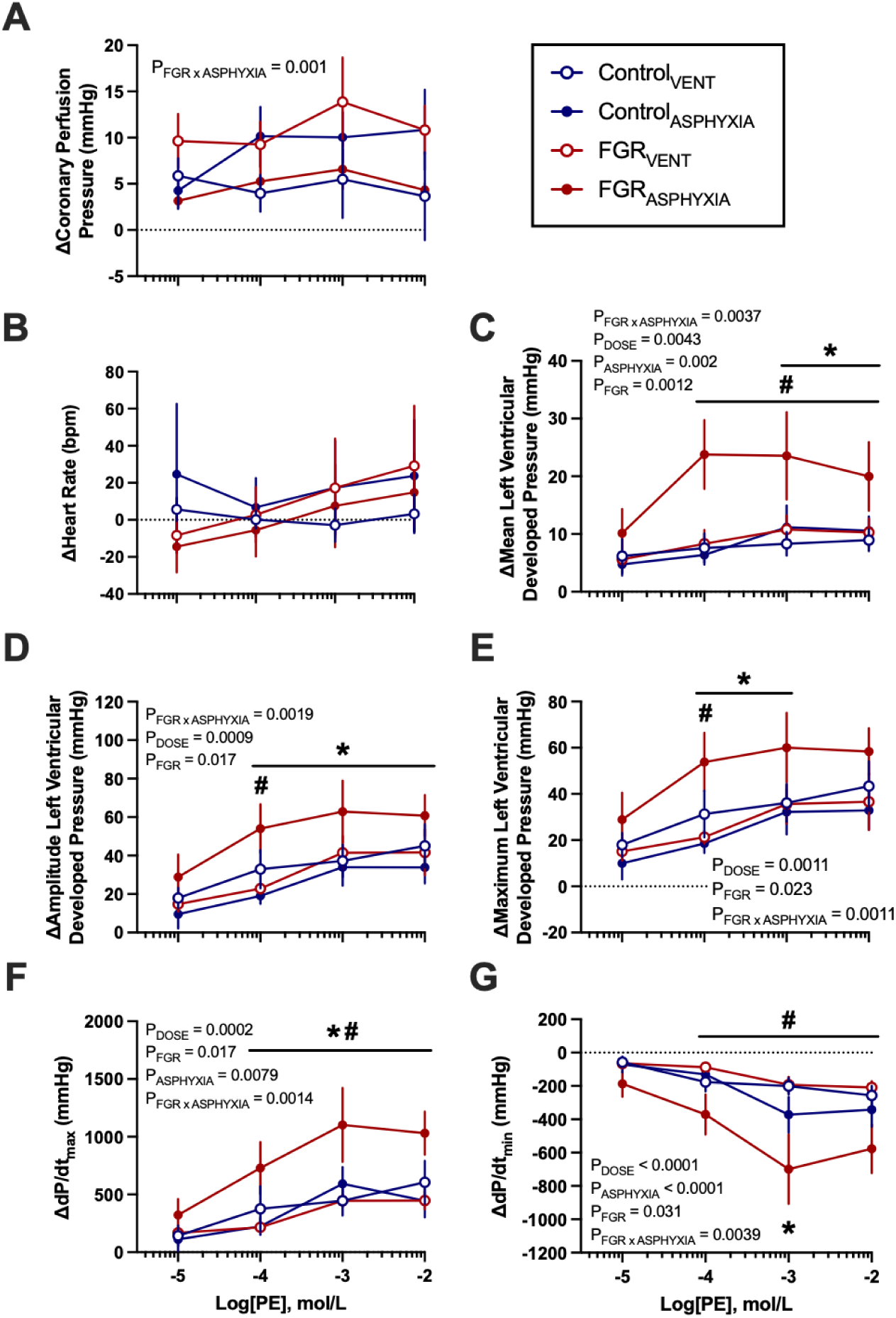
Ex vivo cardiac responses to phenylephrine. Data presented as mean ± standard error of the mean change in (**A**) coronary perfusion pressure, (**B**) heart rate, (**C**) mean left ventricular developed pressure, (**D**) amplitude of left ventricular developed pressure, (**E**) maximum left ventricular developed pressure, (**F**) maximal left ventricular contractility (dP/dt_max_) and (**G**) minimal left ventricular contractility (dP/dt_min_) in response to increasing concentrations of phenylephrine. Groups are ventilated control (Control_VENT_, n=6), ventilated FGR (FGR_VENT_, n=6), asphyxiated control (Control_ASPHYXIA_, n=7), and asphyxiated FGR (FGR_ASPHYXIA_, n=6) lambs. Data analysed via mixed effects analysis with uncorrected Fisher’s least significance multiple comparisons test; statistical significance threshold *p*<0.05. \**p* < 0.05 FGR vs. control; ^#^*p* < 0.05 asphyxia vs. ventilation.

### Histology

Histological assessment of the left ventricle was undertaken to assess the cardiac pathology associated with FGR and asphyxia. Perivascular fibrosis area, but not interstitial fibrosis, was significantly higher in FGR groups compared to controls (p=0.003; Fig. 7A&B). The number of Ki67-positive cells was significantly lower in asphyxia groups (p=0.002; Fig. 7C). Ventricular gross infarct size and left ventricular vascular ɑ_1_ adrenergic receptor content was not different between groups (Supplementary Table 3).

**Figure 7.**
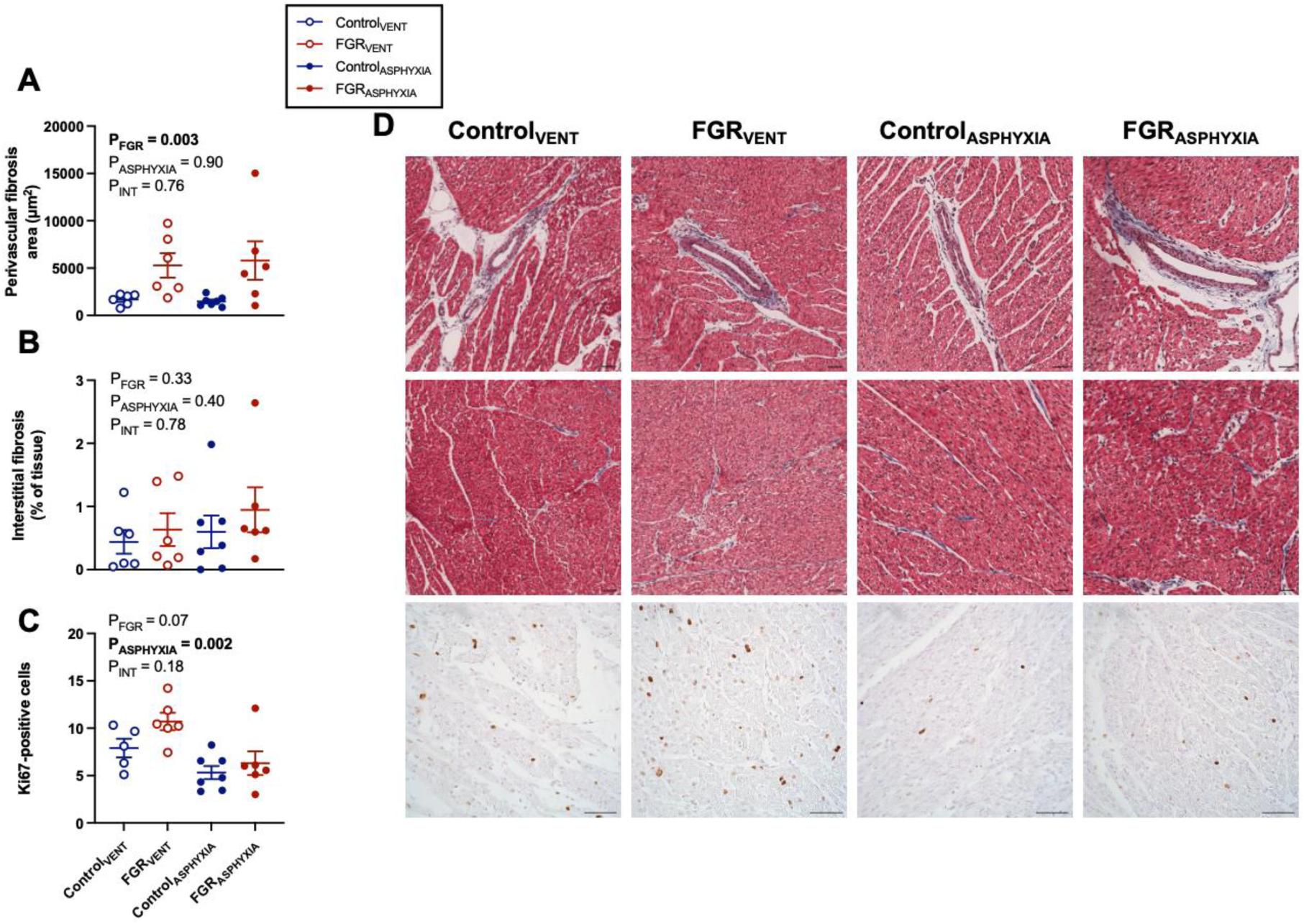
Fibrosis area and cell proliferation in the left ventricle. Data presented as mean ± standard error of (**A**) perivascular fibrosis area, (**B**) % of interstitial fibrosis (relative to tissue area), and (**C**) number of proliferative (Ki67-positive) cells in the left ventricle. (**D**) Representative photomicrographs of Masson’s trichrome-stained left ventricle to evaluate perivascular and interstitial fibrosis, and Ki67-positive cells evaluating cell proliferation. Groups are ventilated control (Control_VENT_, n=6), ventilated FGR (FGR_VENT_, n=6), asphyxiated control (Control_ASPHYXIA_, n=7), and asphyxiated FGR (FGR_ASPHYXIA_, n=6) lambs. Data analysed via two-way ANOVA; statistical significance threshold *p*<0.05. Scale bar = 50µm.

### Gene expression

The mRNA expression of *ɑ_1_-AR*, *β_1_-AR* and *NPPA* in the left ventricle was not different between groups (Supplementary Table 3). The mRNA expression of *NPPB* was higher in FGR_ASPHYXIA_ animals compared to Control_ASPHYXIA_ (p=0.049) and FGR_VENT_ (p=0.03) groups (Supplementary Table 3). The mRNA expression of *SERCA/ATP2A2* was also higher in FGR groups compared to controls (p=0.02; Supplementary Table 3).

## Discussion

FGR and perinatal asphyxia both independently increase the risk of cardiovascular disease^4, 30^, yet their combined effects on cardiovascular function during the perinatal period remain poorly understood. In this current study, we examined the cardiovascular responses of control and FGR lambs to a severe asphyxic insult around the time of birth. We showed that despite FGR lambs having an ability to withstand asphyxia for longer, they had an inability to adequately maintain blood pressure postnatally due to poor vascular contractility. Isolated hearts from FGR lambs demonstrated alterations in α_1_ and β_1_ adrenergic receptor function independent of adrenergic receptor abundance, suggesting potential compensatory mechanisms involving sympathetic activation to maintain contractility during a period of asphyxia. These distinct cardiovascular adaptations highlight how antenatal complications may impact an infant’s response to perinatal asphyxia.

We previously demonstrated that FGR lambs are more resilient to a mild asphyxic event^23^, suggesting that exposure to chronic hypoxia *in utero*, such as in those diagnosed with FGR, results in unique adaptations to an additional asphyxic insult. Here, we have extended these findings by increasing the severity of asphyxia. While a similar resilience to asphyxia is evident in the ability of FGR lambs to withstand asphyxia for a longer duration, there is evidence from both sympathetic dysregulation and cerebral oxygenation that warrants caution over the terminology of resilience to asphyxia. Additionally, in the current study, FGR lambs had a comparable requirement for haemodynamic support and time to ROSC compared to control lambs. The fetal response to profound asphyxia is characterised by a sharp increase in sympathetic activity and subsequent increase in blood pressure, initially activated by α-adrenergic afferents^31^, to maintain perfusion to critical organs through the baroreflex response^21^. In this study, we found that FGR lambs became more hypotensive during asphyxia without a normal corrective change in heart rate, suggesting potential impairments in cardiac baroreflex^32^. Indeed, chronically hypoxaemic fetal sheep at 0.85 gestation had a similar inability to maintain blood pressure during a hypotensive challenge due to an impaired cardiac baroreflex with absent sympathetic regulation^21^. We also investigated whether FGR and control lambs had similar abilities in maintaining blood flow to the brain during asphyxia, given that FGR is associated with the redistribution of cardiac output to the brain^33^. Although carotid blood flow was similar between groups during asphyxia, FGR lambs initially exhibited a stronger redistribution of carotid to femoral blood flow halfway through asphyxia, indicative of a higher output to the cerebral vasculature relative to the periphery. However, this time point also corresponded with an inability of FGR infants to increase their vascular contractility relative to controls. Taken together, these results suggest that vascular adaptations, such as enhanced vasoconstriction, may be driving the increased output to the brain, rather than a substantive increase in cardiac output. Despite this, FGR lambs had lower cerebral oxygenation levels compared to control lambs throughout the entire duration of asphyxia. This highlights potential inefficiencies in oxygen consumption and/or transport despite vascular adaptations to increase oxygenation to the brain. However, little is known about the ability of the FGR brain to efficiently utilise oxygen during acute hypoxia, warranting further investigation in future studies.

In the current study, we found that the isolated hearts of asphyxiated FGR lambs were significantly more contractile in response to α_1_-adrenergic activation compared to asphyxiated controls, suggesting a reliance on α_1_-adrenergic receptor function in the maintenance of autonomic control. This reliance on α_1_ adrenergic receptor function is supported by studies in carunclectomy-induced chronically hypoxic fetal sheep demonstrating a significantly greater hypotensive response to α_1_-adrenergic receptor blockade compared to normoxic fetuses at ∼0.85 gestation^34^. Importantly, the maintenance of blood pressure by α_1_-adrenergic receptor activity was not dependent on post-ganglionic sympathetic activation, signifying an organ-specific mechanism independent of neural stimuli to the adrenal medulla^35^. Despite this, contradicting evidence also exists regarding α_1_-adrenergic receptor modulation in the peripheral vasculature following chronic hypoxia, with sheep studies reporting either impaired^21, 36^ or enhanced^37, 38^ α_1_-adrenergic activity or abundance. Regardless, these data show that there is evolving dysfunction in α_1_-adrenergic function following chronic hypoxia, characterised as a greater reliance of central cardiovascular function (i.e. heart) to maintain cardiac output, with variable effects on the peripheral vasculature. We also found that non-asphyxiated FGR lambs had similar isolated cardiac responses to α_1_-adrenergic receptor activation compared to controls, suggesting that asphyxia independently modulates α-adrenergic activity in FGR hearts to maintain contraction under stress. However, prolonged sympathetic activation can cause cardiomyocyte damage or apoptosis and worsens contractile function, thus increasing the risk of cardiovascular dysfunction in the long-term^39, 40^.

Asphyxiated neonates are at increased risk of haemodynamic instability following the restoration of cardiac output, leading to multi-organ failure and metabolic dysregulation^2^. In the current study, FGR lambs demonstrated a delayed hypertensive overshoot in blood pressure compared to control lambs immediately after ROSC. Rebound hypertension leads to cerebrovascular injury in the neonate^41^. In term lambs, rebound hypertension after ROSC increases blood vessel protein extravasation in the white and grey matter of the brain, which is associated with the loss of tight junction proteins regulating vascular integrity^42^. Preterm FGR infants may be more sensitive to sudden fluctuations in cardiac afterload, leading to an increased risk of cardiovascular and cerebrovascular injury. In particular, low cardiac output in the first 24 hours of life results in increased risk of hypoperfusion-reperfusion brain injury in very low birth weight infants^43^. Fluctuating myocardial perfusion is also likely to cause injury to the FGR heart, as previous studies have shown that isolated hearts from preterm FGR lambs present with larger infarct areas following ischemia-reperfusion injury^44, 45^. This suggests the potential for an increased risk of cardiac injury following a secondary insult in the FGR heart, although this was not evident in the current study where comparable levels of myocardial infarction or fibrosis were observed in FGR and control groups. It is possible that our model either induced a less severe myocardial injury than complete asystole or ischemia-reperfusion injury, or an inadequate duration of exposure to a hypoxic injury. However, asphyxiated FGR lambs had a higher left ventricular mRNA expression of B-type natriuretic peptide, a marker of increased ventricular pressure and wall stress^46^. This suggests that FGR hearts may be more susceptible to subclinical ventricular injury following asphyxia despite having similar infarct sizes to asphyxiated controls. Another important finding was that the return to normotension also occurred faster in FGR lambs. This presents a clinically relevant period where there may be opportunity to therapeutically prevent the overshoot in blood pressure, thereby reducing the risk of rapid fluctuations in blood pressure-associated brain injury.

Impairments in vascular contractility may underlie the altered blood pressure profile observed in FGR lambs during asphyxia and after ROSC. During asphyxia, we observed key differences in the vascular contractility of FGR lambs, including a failure to execute the same sharp increases in the rate of maximal contraction and relaxation in the first quartile of asphyxia. The pulse pressure of FGR lambs was also lower during asphyxia, highlighting a potential risk of cerebral and coronary hypoperfusion as pulse pressure is strongly linked to left ventricular output in neonates^48^. It is likely that poor vascular contractility occurs due to vascular stiffness, as previous preclinical and clinical FGR studies have described early vascular dysfunction and arterial stiffness to occur as early as the perinatal period^36, 47^. Poor vascular contractility also persisted in FGR lambs until 5 minutes after ROSC, highlighting potential impairments in post-resuscitation recovery which may increase the risk of secondary ischemic injury. We investigated whether this vascular stiffness was reflected in myocardial stiffness and found FGR lambs to have an increase in ventricular perivascular fibrosis with no increases to interstitial fibrosis. Excess collagen deposition around the vessels restricts the contractile ability of the myocardium, leading to impairments in systolic and diastolic function^49^. Taken together, poor vascular contractility and fibrosis results in a failure of FGR lambs to dynamically alter and maintain blood pressure after asphyxia, potentially increasing the risk of circulatory instability in the perinatal period.

We also interrogated the responses of the isolated heart to β_1_-adrenergic receptor stimulation, given its importance in maintaining fetal heart rate during brief asphyxic episodes in fetal sheep^31^. Similar to α_1_-adrenergic activation, asphyxiated FGR lambs had a significantly greater response to β_1_-adrenergic stimulation compared to asphyxiated controls. This response was only observed in the lowest dose of dobutamine, suggesting a maximal saturation of β_1_-adrenergic receptors prevented dose-dependent increases in contractility. To date, studies in chronically hypoxic or FGR fetal sheep have only characterised impairments in cardiac function in response to a non-selective β-adrenergic agonist, isoproterenol^44, 45, 50^. We therefore chose to investigate the effects of a clinically relevant β_1_-adrenergic receptor agonist, dobutamine, given its common use in the neonatal intensive care unit to treat hypotensive preterm infants^3^. Activation of β_1_-adrenergic receptors results in the activation of adenylyl cyclase and conversion of ATP into cAMP, ultimately phosphorylating ryanodine receptors and L-type Ca^2+^ channels to allow Ca^2+^ influx into the cytosol. β_1_-adrenergic receptor activation also phosphorylates SERCA/ATP2A2 to enhance calcium reuptake into the sarcoplasmic reticulum, overall resulting in positive inotropy via enhanced calcium cycling. Interestingly, the altered responses to both α_1_ and β_1_ adrenergic activation occurred independently of parenchymal or vascular adrenergic receptor abundance in the left ventricle, and may therefore indicate a greater capacity for calcium handling in the cardiomyocyte. This potential for increased calcium handling is further supported by our finding of increased mRNA expression of SERCA/ATP2A2 in the left ventricles of FGR lambs. Similarly, a study by Lock et al. showed isolated cardiomyocytes from chronically hypoxic mice offspring raised into adulthood had a greater increase in systolic calcium amplitude after isoprenaline stimulation, suggesting sympathetic sensitisation^51^. Taken together, these data suggest that calcium handling may be developmentally programmed during gestation and persist beyond the transitional stage, potentially increasing the risk of sustained sympathetic activation and afterload and increasing the risk of long-term cardiac injury.

There were strengths and limitations to this work. One key strength is our well-defined FGR model of asymmetrical growth restriction, indicated by increased brain-to-body weight ratio with preserved heart-to-body weight ratio, which recapitulates chronic hypoxia and acidemia following placental insufficiency^24^. Our lambs were also delivered preterm at a similar stage of cardiovascular development to a moderately preterm infant^52, 53^ to replicate the large proportion of FGR infants delivered moderately/late preterm to prevent stillbirth^54^. However, we also acknowledge that the investigation of a single gestational age is a limitation given that preterm birth comprises of a broad spectrum of gestational ages and therefore morbidities. Other limitations that may limit the clinical applicability of our findings include the lack of interrogation of intracellular calcium concentrations in isolated cardiomyocytes. We can therefore only extrapolate intracellular calcium handling based on the gene expression of the calcium transporter SERCA/ATP2A2. However, calcium handling also involves other transporters downstream of adrenergic receptor activation, including IP_3_ and ryanodine receptors, which may also provide further insight into the mechanisms underlying the changes in left ventricular contractility. We also acknowledge that our baseline blood gases highlight similar PaO_2_ levels between control and FGR groups, contrary to the characteristic hypoxia traditionally observed in SUAL-induced growth-restricted lambs^24^. This was likely due to the ventilation provided to the ewe during surgical instrumentation of the fetus, increasing placental oxygen transfer prior to collecting the arterial blood sample. This study also lacked an assessment of circulating catecholamine levels, which is known to maintain the chronic redistribution of cardiac output in chronically hypoxaemic fetal sheep and ultimately leads to a blunting of adrenergic receptor activity. However, previous studies have shown no differences in circulating epinephrine or norepinephrine levels in preterm 24-hour old FGR lambs^36^, nor were there differences in the catecholamine levels of asphyxiated preterm FGR and control lambs^23^. Therefore, it is unlikely that an increase in circulating catecholamine levels would have contributed to the altered response to asphyxia.

We have characterised the cardiovascular response between preterm FGR and control lambs to severe perinatal asphyxia by umbilical cord occlusion. We have shown that FGR lambs experience delayed rebound hypertension after resuscitation due to poor myocardial and vascular contractility which may lead to an inadequate perfusion of organs. Importantly, we have identified that these changes occur due to increased ventricular α_1_ and β_1_-adrenergic receptor function independent of receptor abundance. Overall, these data show that FGR infants exposed to asphyxia may be prone to haemodynamic instabilities in the perinatal period, with alterations in sympathetic activation which may persist into the long-term to potentially increase the risk of cardiovascular injury and cardiovascular disease later in life.

## Acknowledgments

We acknowledge the Monash Medical Centre animal house staff and the Monash Animal Research Platform for the care of animals and assistance with experiments.

## Sources of Funding

GRP, BJA, AM and SLM are supported by National Health and Medical Research Council Investigator Grants (funding ID 173731, 1175843, 2008793, 2016688). ZA and BRP were supported by Australian Government Research Training Program Scholarships.

## Disclosures

None.

## Author contributions

ZA, SLM, KJB, GRP and BJA conceived and designed research; ZA, BRP, AES, AT, VZ, YP, IN, MMH, AM, SLM, KJB, GRP and BJA performed experiments; ZA analysed data and prepared figures; ZA, AM, SLM, KJB, GRP and BJA interpreted results of experiments; ZA, KJB, GRP and BJA drafted manuscript; ZA, BRP, AES, AT, VZ, YP, IN, MMH, AM, SLM, KJB, GRP and BJA edited and revised manuscript; ZA, BRP, AES, AT, VZ, YP, IN, MMH, AM, SLM, KJB, GRP and BJA approved final version of manuscript.

## Supplemental Material

Tables S1-3

Figures S1-4

## Notes

### Competing Interest Statement

The authors have declared no competing interest.

## References

1. Lawn J, Shibuya K, Stein C. No cry at birth: Global estimates of intrapartum stillbirths and intrapartum-related neonatal deaths. Bull World Health Organ. 2005;83:409–417

2. Polglase GR, Ong T, Hillman NH. Cardiovascular alterations and multiorgan dysfunction after birth asphyxia. Clin Perinatol. 2016;43:469–483

3. Joynt C, Cheung P-Y. Treating hypotension in preterm neonates with vasoactive medications. Frontiers in Pediatrics. 2018;6

4. Razaz N, Norman M, Alfvén T, Cnattingius S. Low apgar score and asphyxia complications at birth and risk of longer-term cardiovascular disease: A nationwide population-based study of term infants. The Lancet Regional Health – Europe. 2023;24

5. Gurgul S, Buyukakilli B, Komur M, Okuyaz C, Balli E, Ozcan T. Does levetiracetam administration prevent cardiac damage in adulthood rats following neonatal hypoxia/ischemia-induced brain injury? Medicina. 2018;54:12

6. Aziz K, Lee HC, Escobedo MB, Hoover AV, Kamath-Rayne BD, Kapadia VS, et al. Part 5: Neonatal resuscitation: 2020 american heart association guidelines for cardiopulmonary resuscitation and emergency cardiovascular care. Circulation. 2020;142:S524–S550

7. Melamed N, Baschat A, Yinon Y, Athanasiadis A, Mecacci F, Figueras F, et al. Figo (international federation of gynecology and obstetrics) initiative on fetal growth: Best practice advice for screening, diagnosis, and management of fetal growth restriction. International Journal of Gynecology & Obstetrics. 2021;152:3–57

8. Tang L, He G, Liu X, Xu W. Progress in the understanding of the etiology and predictability of fetal growth restriction. Reproduction. 2017;153:R227–R240

9. Malhotra A, Allison BJ, Castillo-Melendez M, Jenkin G, Polglase GR, Miller SL. Neonatal morbidities of fetal growth restriction: Pathophysiology and impact. Front Endocrinol (Lausanne). 2019;10:55

10. Groene SG, Spekman JA, Te Pas AB, Heijmans BT, Haak MC, van Klink JMM, et al. Respiratory distress syndrome and bronchopulmonary dysplasia after fetal growth restriction: Lessons from a natural experiment in identical twins. EClinicalMedicine. 2021;32:100725

11. Suciu LM, Giesinger RE, Mărginean C, Muntean M, Cucerea M, Făgărășan A, et al. Comparative evaluation of echocardiography indices during the transition to extrauterine life between small and appropriate for gestational age infants. Frontiers in Pediatrics. 2023;10

12. Leipälä JA, Boldt T, Turpeinen U, Vuolteenaho O, Fellman V. Cardiac hypertrophy and altered hemodynamic adaptation in growth-restricted preterm infants. Pediatric Research. 2003;53:989–993

13. Sehgal A, Allison BJ, Gwini SM, Miller SL, Polglase GR. Cardiac morphology and function in preterm growth restricted infants: Relevance for clinical sequelae. J Pediatr. 2017;188:128–134.e122

14. Cohen E, Wong FY, Wallace EM, Mockler JC, Odoi A, Hollis S, et al. Fetal-growth-restricted preterm infants display compromised autonomic cardiovascular control on the first postnatal day but not during infancy. Pediatric Research. 2017;82:474–482

15. Wood S, Crawford S, Hicks M, Mohammad K. Hospital-related, maternal, and fetal risk factors for neonatal asphyxia and moderate or severe hypoxic-ischemic encephalopathy: A retrospective cohort study. The journal of maternal-fetal & neonatal medicine. 2021;34:1448–1453

16. Liu J, Wang XF, Wang Y, Wang HW, Liu Y. The incidence rate, high-risk factors, and short- and long-term adverse outcomes of fetal growth restriction: A report from mainland china. Medicine (Baltimore). 2014;93:e210

17. Piscopo BR, Malhotra A, Hunt RW, Davies-Tuck ML, Palmer KR, Sutherland AE, et al. The interplay between birth weight and intraventricular hemorrhage in very preterm neonates—a retrospective cohort study. American Journal of Obstetrics & Gynecology MFM. 2025;7:101628

18. Low JA, Boston RW, Pancham SR. Fetal asphyxia during the intrapartum period in intrauterine growth-retarded infants. Am J Obstet Gynecol. 1972;113:351–357

19. Giussani DA. The fetal brain sparing response to hypoxia: Physiological mechanisms. J Physiol. 2016;594:1215–1230

20. Ahmadzadeh E, Polglase GR, Stojanovska V, Herlenius E, Walker DW, Miller SL, et al. Does fetal growth restriction induce neuropathology within the developing brainstem? J Physiol. 2023;601:4667–4689

21. Allison BJ, Brain KL, Niu Y, Kane AD, Herrera EA, Thakor AS, et al. Altered cardiovascular defense to hypotensive stress in the chronically hypoxic fetus. Hypertension. 2020;76:1195–1207

22. Lear CA, Georgieva A, Beacom MJ, Wassink G, Dhillon SK, Lear BA, et al. Fetal heart rate responses in chronic hypoxaemia with superimposed repeated hypoxaemia consistent with early labour: A controlled study in fetal sheep. BJOG: An International Journal of Obstetrics & Gynaecology. 2023;130:881–890

23. Oyang M, Piscopo BR, Zahra V, Malhotra A, Sutherland AE, Sehgal A, et al. Cardiovascular responses to mild perinatal asphyxia in growth-restricted preterm lambs. American Journal of Physiology-Heart and Circulatory Physiology. 2023;325:H1081–H1087

24. Sutherland AE, White TA, Rock CR, Piscopo BR, Dudink I, Inocencio IM, et al. Phenotype of early-onset fetal growth restriction in sheep. Frontiers in Endocrinology. 2024;15

25. Alves de Alencar Rocha AK, Allison BJ, Yawno T, Polglase GR, Sutherland AE, Malhotra A, et al. Early-versus late-onset fetal growth restriction differentially affects the development of the fetal sheep brain. Dev Neurosci. 2017;39:141–155

26. de Jager J, Pothof R, Crossley KJ, Schmölzer GM, Te Pas AB, Galinsky R, et al. Evaluating the efficacy of endotracheal and intranasal epinephrine administration in severely asphyxic bradycardic newborn lambs: A randomised preclinical study. Arch Dis Child Fetal Neonatal Ed. 2025;110:207–212

27. Mulrooney N, Champion Z, Moss TJ, Nitsos I, Ikegami M, Jobe AH. Surfactant and physiologic responses of preterm lambs to continuous positive airway pressure. Am J Respir Crit Care Med. 2005;171:488–493

28. Hess RM, Niu Y, Garrud TAC, Botting KJ, Ford SG, Giussani DA. Embryonic cardioprotection by hydrogen sulphide: Studies of isolated cardiac function and ischaemia-reperfusion injury in the chicken embryo. J Physiol. 2020;598:4197–4208

29. Hansen TS, Karimi Galougahi K, Tang O, Tsang M, Scherrer-Crosbie M, Arystarkhova E, et al. The fxyd1 protein plays a protective role against pulmonary hypertension and arterial remodeling via redox and inflammatory mechanisms. American Journal of Physiology-Heart and Circulatory Physiology. 2023;326:H623–H635

30. Crispi F, Bijnens B, Figueras F, Bartrons J, Eixarch E, Le Noble F, et al. Fetal growth restriction results in remodeled and less efficient hearts in children. Circulation. 2010;121:2427–2436

31. Galinsky R, Lear CA, Yamaguchi K, Wassink G, Westgate JA, Bennet L, et al. Cholinergic and β-adrenergic control of cardiovascular reflex responses to brief repeated asphyxia in term-equivalent fetal sheep. American Journal of Physiology-Regulatory, Integrative and Comparative Physiology. 2016;311:R949–R956

32. Andriessen P, Oetomo SB, Peters C, Vermeulen B, Wijn PF, Blanco CE. Baroreceptor reflex sensitivity in human neonates: The effect of postmenstrual age. J Physiol. 2005;568:333–341

33. Miller SL, Huppi PS, Mallard C. The consequences of fetal growth restriction on brain structure and neurodevelopmental outcome. J Physiol. 2016;594:807–823

34. Danielson L, McMillen IC, Dyer JL, Morrison JL. Restriction of placental growth results in greater hypotensive response to alpha-adrenergic blockade in fetal sheep during late gestation. J Physiol. 2005;563:611–620

35. Darby JRT, Varcoe TJ, Holman SL, McMillen IC, Morrison JL. The reliance on α-adrenergic receptor stimuli for blood pressure regulation in the chronically hypoxaemic fetus is not dependent on post-ganglionic activation. J Physiol. 2021;599:1307–1318

36. Rock CR, White TA, Piscopo BR, Sutherland AE, Pham Y, Camm EJ, et al. Cardiovascular decline in offspring during the perinatal period in an ovine model of fetal growth restriction. Am J Physiol Heart Circ Physiol. 2023;325:H1266–h1278

37. Brain KL, Allison BJ, Niu Y, Cross CM, Itani N, Kane AD, et al. Intervention against hypertension in the next generation programmed by developmental hypoxia. PLoS Biol. 2019;17:e2006552

38. Herrera EA, Rojas RT, Krause BJ, Ebensperger G, Reyes RV, Giussani DA, et al. Cardiovascular function in term fetal sheep conceived, gestated and studied in the hypobaric hypoxia of the andean altiplano. J Physiol. 2016;594:1231–1245

39. Gronda E, Dusi V, D’Elia E, Iacoviello M, Benvenuto E, Vanoli E. Sympathetic activation in heart failure European Heart Journal Supplements. 2022;24:E4–E11

40. Borovac JA, D’Amario D, Bozic J, Glavas D. Sympathetic nervous system activation and heart failure: Current state of evidence and the pathophysiology in the light of novel biomarkers. World J Cardiol. 2020;12:373–408

41. Badurdeen S, Roberts C, Blank D, Miller S, Stojanovska V, Davis P, et al. Haemodynamic instability and brain injury in neonates exposed to hypoxia– ischaemia. Brain Sciences. 2019;9:49

42. Polglase GR, Blank DA, Barton SK, Miller SL, Stojanovska V, Kluckow M, et al. Physiologically based cord clamping stabilises cardiac output and reduces cerebrovascular injury in asphyxiated near-term lambs. Archives of Disease in Childhood - Fetal and Neonatal Edition. 2018;103:F530–F538

43. Kluckow M, Evans N. Low superior vena cava flow and intraventricular haemorrhage in preterm infants. Arch Dis Child Fetal Neonatal Ed. 2000;82:F188–194

44. Tare M, Miller SL, Wallace EM, Sutherland AE, Yawno T, Coleman HA, et al. Glucocorticoid treatment does not alter early cardiac adaptations to growth restriction in preterm sheep fetuses: Growth-restricted heart not worsened by antenatal glucocorticoid. BJOG. 2012;119:906–914

45. Tare M, Parkington HC, Wallace EM, Sutherland AE, Lim R, Yawno T, et al. Maternal melatonin administration mitigates coronary stiffness and endothelial dysfunction, and improves heart resilience to insult in growth restricted lambs. J Physiol. 2014;592:2695–2709

46. Alter P, Rupp H, Rominger MB, Vollrath A, Czerny F, Klose KJ, et al. Relation of b-type natriuretic peptide to left ventricular wall stress as assessed by cardiac magnetic resonance imaging in patients with dilated cardiomyopathy. Can J Physiol Pharmacol. 2007;85:790–799

47. Sehgal A, Allison BJ, Gwini SM, Menahem S, Miller SL, Polglase GR. Vascular aging and cardiac maladaptation in growth-restricted preterm infants. J Perinatol. 2018;38:92–97

48. Kharrat A, Rios DI, Weisz DE, Giesinger RE, Groves A, Yang J, et al. The relationship between blood pressure parameters and left ventricular output in neonates. J Perinatol. 2019;39:619–625

49. de Boer RA, De Keulenaer G, Bauersachs J, Brutsaert D, Cleland JG, Diez J, et al. Towards better definition, quantification and treatment of fibrosis in heart failure. A scientific roadmap by the committee of translational research of the heart failure association (hfa) of the european society of cardiology. Eur J Heart Fail. 2019;21:272–285

50. Browne VA, Stiffel VM, Pearce WJ, Longo LD, Gilbert RD. Cardiac beta-adrenergic receptor function in fetal sheep exposed to long-term high-altitude hypoxemia. Am J Physiol. 1997;273:R2022–2031

51. Lock MC, Patey OV, Smith KLM, Niu Y, Jaggs B, Trafford AW, et al. Maladaptive cardiomyocyte calcium handling in adult offspring of hypoxic pregnancy: Protection by antenatal maternal melatonin. J Physiol. 2024;602:6683–6703

52. Burrell JH, Boyn AM, Kumarasamy V, Hsieh A, Head SI, Lumbers ER. Growth and maturation of cardiac myocytes in fetal sheep in the second half of gestation. The Anatomical Record Part A: Discoveries in Molecular, Cellular, and Evolutionary Biology. 2003;274A:952–961

53. Jonker SS, Zhang L, Louey S, Giraud GD, Thornburg KL, Faber JJ. Myocyte enlargement, differentiation, and proliferation kinetics in the fetal sheep heart. J Appl Physiol (1985). 2007;102:1130–1142

54. Rabinovich A, Tsemach T, Novack L, Mazor M, Rafaeli-Yehudai T, Staretz-Chacham O, et al. Late preterm and early term: When to induce a growth restricted fetus? A population-based study. J Matern Fetal Neonatal Med. 2018;31:926–932

